# Mucin MUC5B levels are reduced in the colonic epithelia of ulcerative colitis patients

**DOI:** 10.1101/2024.02.21.581373

**Authors:** Wei Ding, Walter A. Koltun, Gregory S. Yochum

## Abstract

**Background & Aims:** Ulcerative colitis (UC) is an inflammatory bowel disease (IBD) where a defect in the colonic epithelial barrier elicits a chronic and unresolved inflammatory response against luminal microbiota. Mucins are heavily glycosylated proteins secreted by goblet cells that form gelatinous layers to protect the colonic epithelium. Whereas the role of the major colonic mucin, MUC2, is established in UC, few reports have analyzed MUC5B in this disease. In this study, we investigated MUC5B expression in colonic tissues and organoid cultures from UC patients and non-IBD controls.

**Methods:** *MUC5B* transcripts were analyzed in colonic regions from UC patients and controls using RT-qPCR. MUC5B protein levels in colonic tissues were analyzed and quantified by immunohistochemistry. Patient-derived epithelial organoids were stained for MUC5B, E-cadherin and F-actin. Organoids were treated with interleukin 1 beta (IL-1β) and *MUC5B* transcripts were analyzed by RT-qPCR. MUC5B proteins were assessed in IL-1β-treated organoids by immunofluorescent microscopy.

**Results:** *MUC5B* transcripts and proteins were expressed in the healthy colon and their levels were decreased in UC tissues. MUC5B proteins were reduced in organoids derived from UC tissues. IL-1β treatment of control organoids stimulated expression of acidic mucins, *MUC5B* transcripts and MUC5B proteins. *MUC5B* transcripts were not induced by IL-1β in UC-derived organoids.

**Conclusions:** MUC5B levels are reduced in UC colons and its expression is induced by the IL-1β pro-inflammatory cytokine. These results suggest that MUC5B contributes to the protective colonic mucin layer and that this function is likely compromised in UC.

## INTRODUCTION

Ulcerative colitis (UC) is one of two major types of inflammatory bowel disease (IBD) where chronic inflammation deteriorates the colonic epithelium (1). UC affects the colon with a continuous disease presentation and inflammation in UC primarily involves the mucosal layer (1). It is thought that UC develops in genetically susceptible individuals that are incapable of fully resolving an inflammatory response against commensal or pathogenic microbiota in the intestinal lumen (2).

A single layer of epithelial cells lines the intestines and provides a physical barrier that prevents inappropriate penetration of bacteria into underlying tissues (3). In the small intestine, the epithelium is arranged into villi that protrude into the lumen and crypts that extend into the mucosal layer (4). In the colon, the villi are absent but crypt invaginations are maintained (4). Goblet cells, found in the differentiated epithelium, secrete heavily-glycosylated mucin proteins that form a gelatinous layer which protects the epithelium from mechanical stress and direct contact with the luminal microbiota (5). A single, tightly packed mucinous layer is found in the small intestines whereas a mucous bilayer lines the colons with a more-dense epithelial proximal lattice coated with a looser gel coating that is impregnated by commensal bacteria (5, 6). Transmembrane mucins are physically-anchored to the cell membrane whereas secreted, or gel-forming, mucins form large polymers (5, 6).

Two gel-forming mucins expressed in the intestines are MUC2 and MUC5B (6, 7). Of these, MUC2 plays a major role in protecting the colonic epithelium as *Muc2^-/-^* mice develop colitic phenotypes including weight loss, a deteriorated intestinal epithelial architecture, infiltration of immune cells into the mucosal layer and elevated expression of pro-inflammatory cytokines (8). Moreover, goblet cell populations and MUC2 levels are depleted in colonic segments derived from UC patients (9, 10). MUC5B is expressed in colonic goblet cells, but is not expressed in the small intestines, suggesting that it may play a tissue-specific role in protecting the colonic epithelium (11). In line with this possibility, we found in previous transcriptome analyses of colonic tissues from UC patients that increased *MUC5B* expression correlated with delayed requirement for surgical intervention in a subset of these patients (12). In addition, Vancamelbeke et al. reported that *MUC5B* transcripts were up-regulated in the inflamed colons of patients with active UC versus controls (13). However, a recent study found no significant difference in MUC5B protein levels in colonic biopsies of controls versus UC tissues with active disease or UC tissues during disease remission (14). These differing results indicate that the role of MUC5B in the pathophysiology of UC is not fully understood.

IL-1β is a proinflammatory cytokine whose expression is increased following recognition of bacterial products through intracellular pattern recognition receptors and activation of the inflammasome (15). IL-1β recruits and activates immune cells including macrophages, dendritic cells and neutrophils to the site of inflammation (15) and higher levels of IL-1β are found in the intestinal mucosa of IBD patients (16, 17). In addition, IL-1β has been shown to increase intestinal epithelial tight junction permeability through multiple mechanisms (18).

In this study, we evaluated MUC5B expression in healthy colonic segments from non-IBD controls (hereafter referred to as controls) and in inflamed colonic segments from UC patients. We found that MUC5B transcripts and proteins were decreased in UC patient tissues and that these phenotypes were maintained in colonic organoid cultures derived from these tissues. Treating healthy colonic organoids with IL-1β stimulated expression of MUC5B; however, IL-1β treatment failed to induce *MUC5B* transcripts in organoids derived from UC patients. Together, these results suggest that MUC5B contributes to the protective colonic mucin bilayer and that this function is compromised in UC patients.

## MATERIALS and METHODS

### Patient samples

Patients were consented to have surgically-resected intestinal tissues stored in the Carlino Family Inflammatory Bowel and Colorectal Disease (IBCRD) biobank at the Pennsylvania State University College of Medicine. This study, and the IBCRD biobank, were approved by the Pennsylvania State University College of Medicine Institutional Review Board (PRAMSHY98-057). Full-thickness intestinal tissue segments were obtained from UC patients and controls.

Controls were non-IBD patients that underwent surgery for colorectal cancer, volvulus, slow-transit constipation or other digestive disorders. Healthy colonic tissues from these patients were evaluated. Immediately after the tissues were resected, the operating surgeon transported them to the gross pathology lab where they were collected by a technician and transported to the IBCRD biobank lab on ice. The tissues were then separated into ∼100 mg aliquots that were stored in formalin, RNA-later (ThermoFisher, AM7021) or flash-frozen in liquid nitrogen. Samples in RNA-later, or that were flash-frozen, were stored in a −80°C freezer until use for experiments.

### RT-qPCR

RNA-later was removed and the tissue samples were resuspended in 0.5 ml of TRIzol reagent (Invitrogen, ThermoFisher, 15596018), homogenized using a disposable plastic micropestle and total RNA was isolated following the manufacturer’s instruction. RNAs were further purified using the RNAEasy Mini Kit (Qiagen, 74104). RNAs were quantified and their integrities were assessed using a Nanodrop bioanalyzer (Agilent). From 400 ng of total RNA, cDNAs were synthesized using random hexamer primers and the Versco cDNA kit (ThermoFisher, AB1453B) following the manufacturer’s instructions. The cDNAs were diluted to 20 ng/μl and 5 μl was subjected to quantitative PCR using Taqman probes in 10 μl reactions using the Gene Expression Master Mix kit (ThermoFisher, 4369016). The following Taqman probes were purchased from ThermoFisher: *MUC5B*, Hs00861588_m1; *MUC2*, Hs00894041_g1; *VIL1* Hs01031739_m1; *CDH1*, Hs01023894_m1; *GAPDH*, Hs02758991_g1. Reactions were incubated at 95°C for 2 min, followed by 60 cycles at 95°C for 20s, 60°C for 20 s and 72°C for 30 s. Data is presented as relative transcript levels using the 2^-△△CT^ method after normalizing to *GAPDH* or *CDH1*.

### Immunohistochemical analyses of patient samples

Full-thickness colonic tissues fixed in formalin were embedded in paraffin blocks and sectioned onto slides using a cryostat. Slides were rehydrated and subjected to antigen retrieval by incubating them in a rice steamer containing 10 mM sodium citrate (Na_3_C_6_H_5_O_7_), 0.05% tween-20, pH=6.0 for 20 min. Peroxidase was quenched by incubating them in 3% hydrogen peroxide (H_2_O_2_) for 10 min and they were then blocked with 1% normal horse serum (Vector Laboratories, SP-2001) at room temperature for 15 min. The slides were incubated with anti-MUC2 antibodies (1:500, Santa Cruz, Ccp58, sc-7314) or anti-MUC5B antibodies (1:200, Santa Cruz, 5B#-19-2E, sc-21768) overnight at 4°C. Afterwards, the slides were incubated with horse biotinylated anti-mouse/ rabbit IgG ABC-HRP peroxidase at room temperature for 1 hr using the VECTASTAIN Elite ABC Universal Kit (Vector Laboratories, PK-6200). The slides were then washed with 1X PBS for 10 min at room temperature and incubated with VECTASTATIN Elite ABC reagent (Vector Laboratories, PK-6200) for 30 min at room temperature. The slides were developed using DAB liquid chromogen (Vector Laboratories, SK-4100) and were counterstained with hematoxylin before mounting. Slides were visualized using a microscope (Olympus, BH-2), and images were collected using MU500 camera (AmScope).

### Organoid cultures

We followed guidelines from Stem Cell Technologies for culturing human colonic organoids with some modifications. Briefly, the mucosal layer was removed from an approximately 1 cm^3^ fresh section of human colonic tissue and washed twice with 1X Dulbecco’s phosphate-buffered saline (DPBS, ThermoFisher, 14040133). The mucosa was transferred to a 5-ml conical tube containing 0.5 ml crypt isolation buffer (2 mM EDTA, pH=8.0; 43.4 mM sucrose; 54.9 mM D-sorbitol dissolved in 1X DPBS) and minced with a sterile scissor into approximately 0.5 mm^3^ fragments. The mixture was transferred to a 15-ml conical tube containing 10 ml of crypt isolation buffer and 0.5 ml of a 100 X penicillin/streptomycin antibiotic stock solution (Corning, 30-002-CI). The tubes were incubated on a rocking platform at 40 rpm for 30 min on ice, centrifuged at 300 X g for 5 min and the pellets were resuspended in 2 ml basal medium [DMEM/F12, (Gibco, 11330-032) supplemented with 1% bovine serum albumin (Gemini, 700-100p)]. Crypts were isolated by triturating the mixture for 20 repetitions, passing them through a 100 μm strainer (VWR, 76327-102) and then centrifuging them at 300 X g for 5 min. Pellets were resuspended in 50 μl basal medium, an equal volume of pre-chilled Matrigel (Corning, 356231) was added and 50 μl of the mixture was placed in the center of a well in a 24-well cell culture plate. The plates were incubated at 37°C for 20 min to allow the domes to solidify after which 750 μl of human intestinal organoid growth media (Human IntestiCult Organoid Growth Medium, hIOGM, StemCell Technologies, 06010) was added along with 10 μM ROCK inhibitor (Y-27632, Miltenyi, 130-106-138).

### Indirect-immunofluorescence staining of organoid cultures

Organoids cultured on circular coverslips were rinsed with 1X DPBS lacking calcium and magnesium (Corning, 21-031-CV) and fixed with 4% paraformaldehyde at room temperature for 30 min. The coverslips were washed three times with 1X DPBS and incubated for 2 hrs. at room temperature in blocking/staining buffer (5% donkey serum, 0.2% Triton X-100, 1X DPBS). The cover slips were then incubated with the following primary antibodies diluted in blocking/staining buffer overnight at 4°C: anti-MUC5B (1:200, Atlas Antibodies, HPA008246); anti-E-cadherin (1:200, Cell Signaling, 24E10, 3195); anti-F-actin (1:500, Phalloidin-iFluor 647, Abcam, ab176759). The next day, coverslips were washed three times for 10 min and then incubated with the appropriate secondary antibodies (1:500, Alexa donkey anti-rabbit 568, Invitrogen, A10042, or Alexa donkey anti-mouse 488, Invitrogen, A21202) for two hrs at room temperature. The organoids were washed three times in 1X DPBS, for 10 min each, and stained with 4’,6-diamdino-2-phenylindole (DAPI) (1:500, ThermoFisher, 62248) in 1X DPBS for 20 min. After three, 5 min washes in 1X DPBS, the coverslips were inverted onto a glass slide, and treated with prolong anti-fade mounting solution (Invitrogen, P36982). Images were analyzed using a Leica SP8 confocal microscope.

### Organoid treatment with IL-1β

Organoids were differentiated in the IntestiCult Organoid Differentiation Medium (StemCell Technologies, 100-0214) supplemented with 5 μM DAPT ((2S)-N-[(3,5-difluorophenyl) acetyl-L-alanyl-2-phenyl] glycine 1,1-dimethylethyl ester, Tocris, 2634) for 48 hrs. The differentiated organoids were treated with the indicated doses of recombinant human IL-1β (R&D system, 201-LB) for an additional 48 hrs.

### Alcian blue staining

Organoids were fixed with 4% paraformaldehyde and 0.1% glutaraldehyde in 1X DPBS for 30 min at room temperature and then rinsed in 1X DPBS three times for 10 min each. The organoid domes were transferred to a 50-ml conical tube containing a 20% sucrose/ 1X DPBS solution and incubated at 4°C until the organoids settled to the bottom of the tube. The organoids were transferred to an embedding mold containing optimal cutting temperature (OCT) compound (Tissue-Tek, 4583), snap frozen in liquid nitrogen and stored at −80°C. The organoids blocks were cut as 10 µm-thickness sections onto slides using a cryotome, the slides were air-dried and then rinsed three times with distilled water. The slides were incubated in 3% acetic acid for three min at room temperature, stained with 1% Alcian blue (pH 2.5) solution (ACROS organics, 400460250) for 30 min at room temperature, washed under running water for 10 min and then rinsed in distilled water. The slides were then counterstained in nuclear fast red (Poly Scientific R&D corporation, s248-8OZ) for 5 min at room temperature, washed three times in distilled water, mounted with a coverslip and sealed with Cytoseal 60 (Epredia, 8310-4).

### Statistics

An unpaired two-tailed Student’s t-test was used in experiments comparing results between two groups. One-way ANOVA analyses were used in experiments comparing results between multiple groups. A *P*-value of less than 0.05 was considered a statistically significant difference in all cases.

## RESULTS

### MUC5B is expressed in healthy colons

Earlier studies reported that MUC5B is expressed at the transcript and protein levels in the colon (11, 12). To confirm these findings in an independent cohort, we isolated RNAs from full-thickness and healthy colonic segments of control patients that underwent surgery for non-IBD diseases at the Pennsylvania State Milton S. Hershey Medical Center. We included segments from the ascending, descending and sigmoid colons from five independent patients for each.

After converting the RNAs to cDNAs, we conducted real-time quantitative PCR analysis to measure transcripts. In addition to *MUC5B*, we surveyed *MUC2* transcripts as a positive control and *VILLIN* transcripts as an internal control. Whereas we readily detected *MUC5B* transcripts in each colonic segment, their levels were lower than *MUC2* (Fig. 1A). Through immunohistochemical analysis, we found that MUC5B proteins were expressed in colonic epithelial cells, although at qualitatively lower levels than MUC2 (Fig. 1B).

**Fig. 1.**
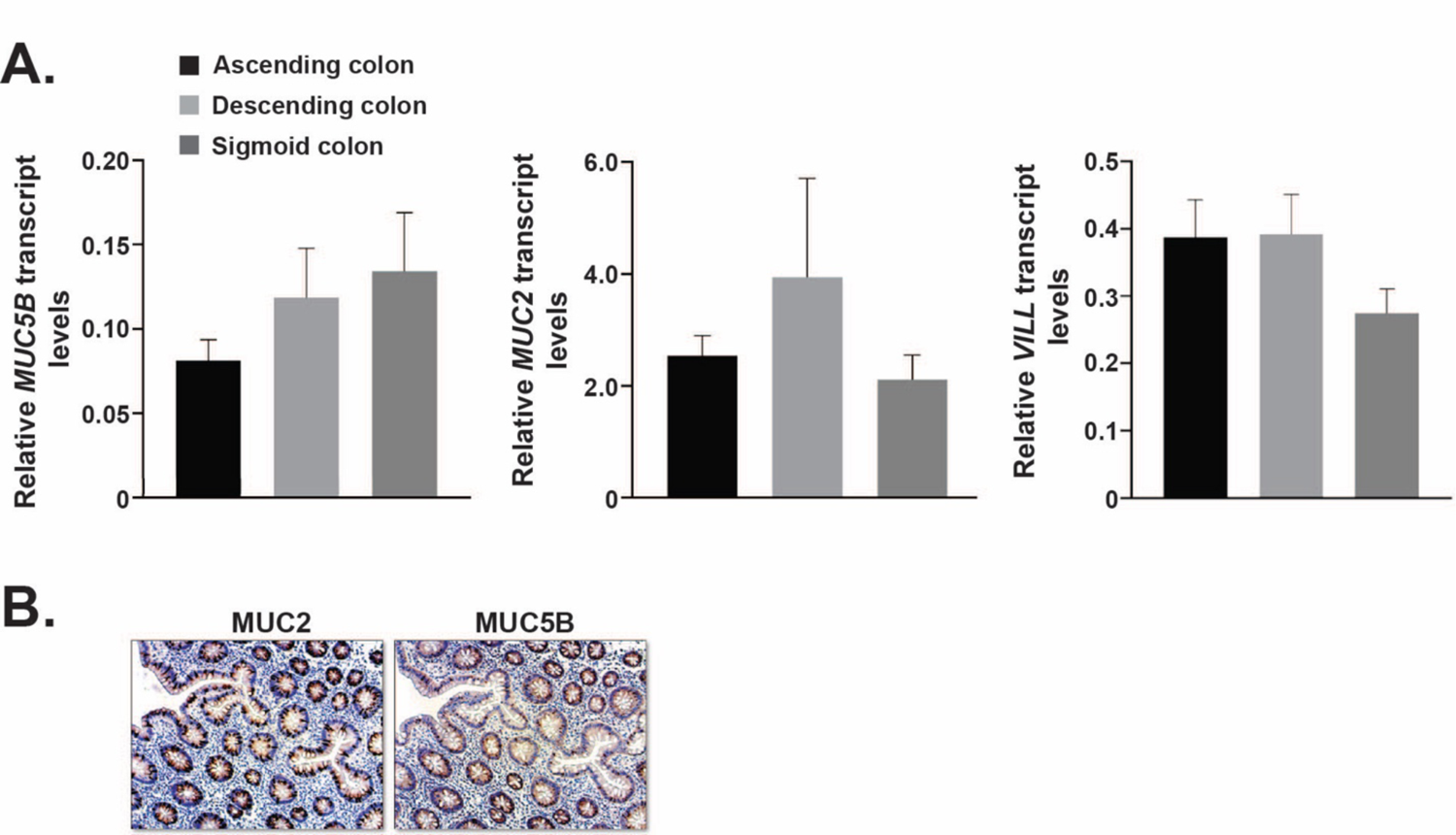
MUC5B is expressed in healthy colons. (A) RT-qPCR analyses of *MUC2*, *MUC5B*, and *VILLIN* (*VILL*) transcripts in RNAs purified from segments of ascending, descending and sigmoid colons. Healthy colonic tissue sections from non-IBD controls (n=5 for each region) were evaluated. Expression levels were normalized to E-cadherin (*CDH1*). Error bars represent SEM. (B) Immunohistochemical analysis of MUC2 and MUC5B in control colonic segments.

### MUC5B is reduced in inflamed colons from UC patients

To evaluate MUC5B expression in colonic segments from UC patients, we established a scoring system based on staining intensity where sections were assigned a score of 0-3, with 0 representing little to no staining and 3 representing robust staining (Fig. 2A). A representative image of an inflamed UC colonic segment with little staining is shown in Fig. 2B. Expanding this analysis to include additional controls (n=13) and inflamed UC segments (n=28) found that MUC5B protein levels are reduced in the colonic epithelia of UC patients (Fig. 2C, Suppl. Table I).

**Fig. 2.**
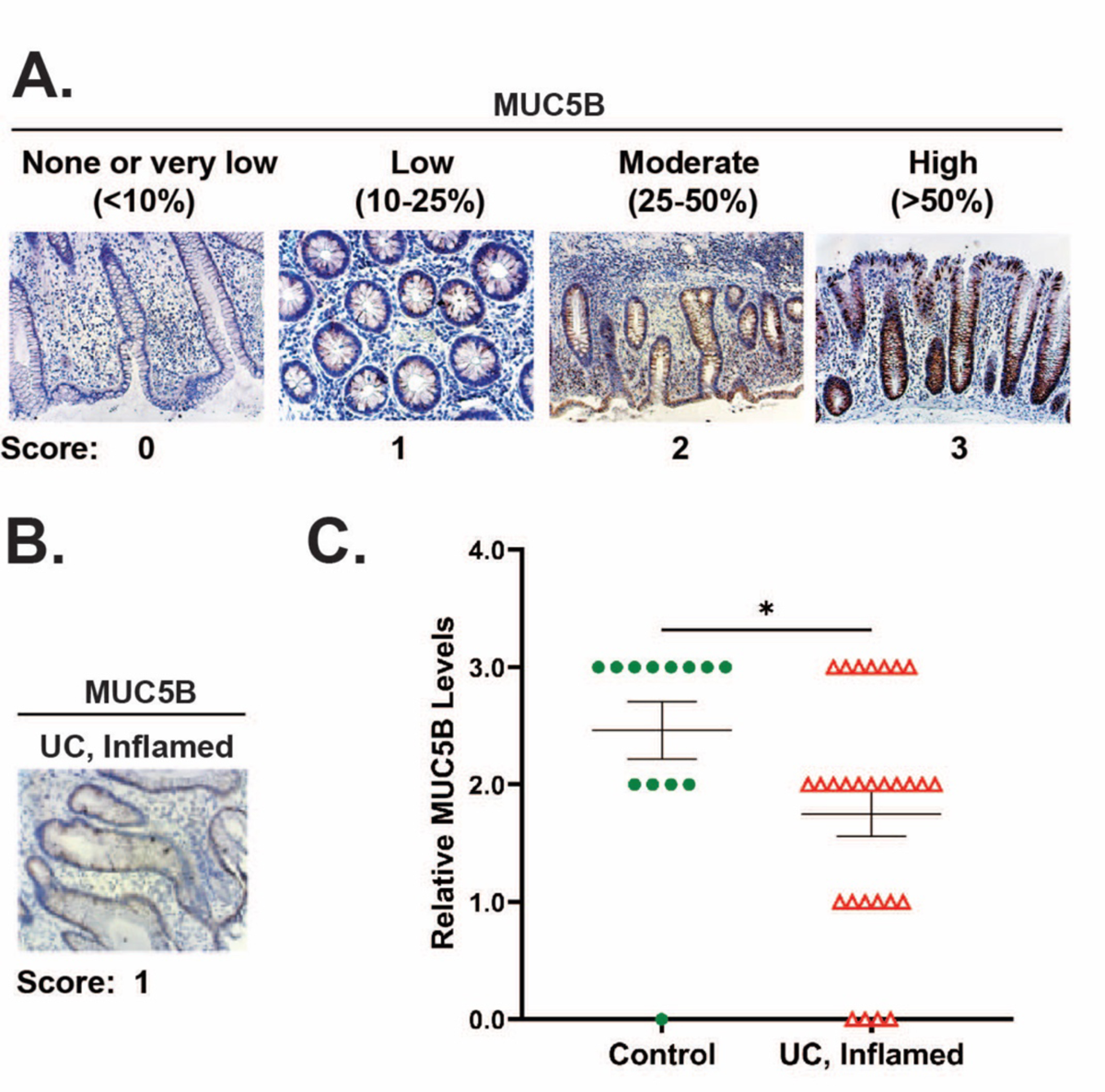
MUC5B is reduced in inflamed colons from UC patients. (A) Scoring system to analyze MUC5B proteins in intestinal tissues. Positively stained cells were quantified per field of view under 10X magnification. At least five fields of view were assessed for each tissue and staining patterns were scored as follows: 0 (negative or very low numbers of positive cells, 0-10%), 1 (low, 10-25%), 2 (moderate, 25-50%) or 3 (high, >50%). (B) Immunohistochemical analysis of MUC5B in an inflamed colonic segment from a UC patient. (C) Quantification of MUC5B staining in colons from non-IBD controls (n=12) and in inflamed colons from UC patients (n=28). Error bars represent SEM, \**P* < 0.05.

### MUC5B is reduced in colon organoid cultures from UC patients

Colonic organoids are three-dimensional cultures derived from the epithelium that can be propagated *in vitro* (19, 20). To determine whether the reduced MUC5B phenotype in UC patient colons is maintained in organoids, we established independent cultures from non-IBD control (n=3) and inflamed UC (n=3) colons. After culturing the organoids for three passages, we differentiated them for 48 hrs. and subsequently analyzed *MUC5B* transcripts. In comparison to control cultures, UC organoids displayed a 60% decrease in *MUC5B* expression (Fig. 3A). To determine whether this decrease in transcripts was also seen at the protein level, we conducted immunofluorescent staining analysis and found decreased MUC5B in UC colonic organoids versus controls (Fig. 3B).

**Fig. 3.**
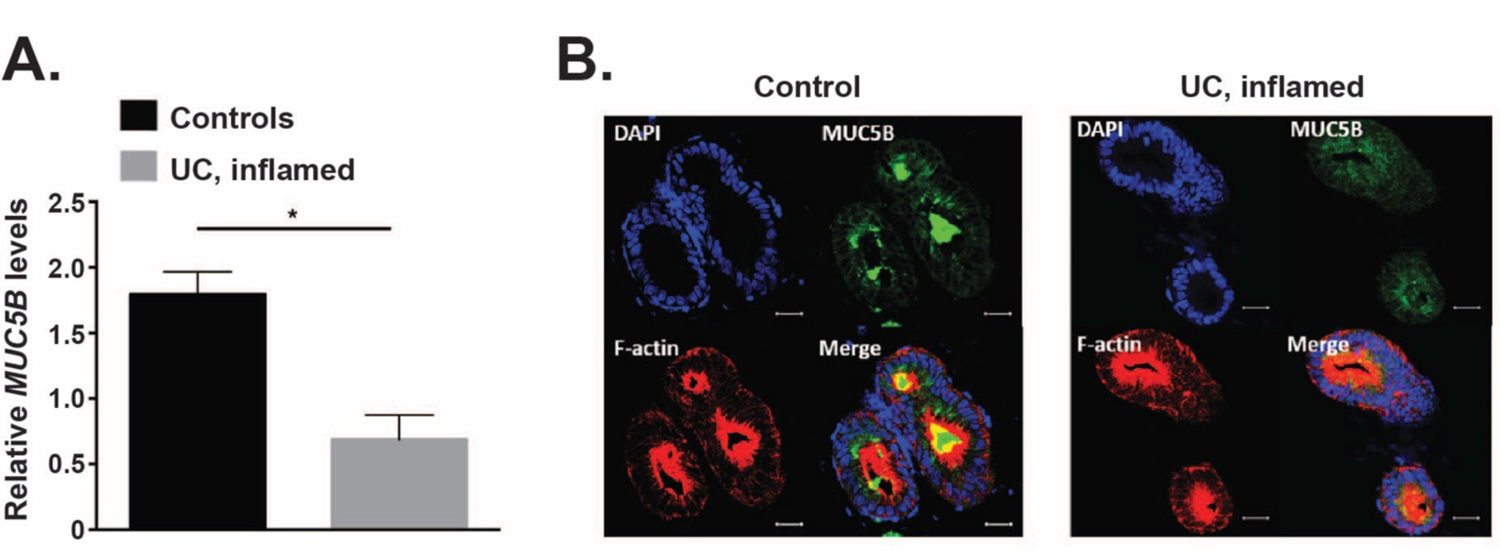
MUC5B is reduced in colon organoid cultures from UC patients. (A) RT-qPCR analysis of *MUC5B* transcript levels in colonic organoid cultures derived from control (n=3) and inflamed colonic segments from UC patients (n=3). *MUC5B* levels are normalized to *GAPDH*. Error bars are SEM, \**P* < 0.05. (B) Colonic organoid cultures derived from control and inflamed UC colonic segments that were stained with DAPI to detect cell nuclei, anti-F-actin antibodies or anti-MUC5B antibodies. Scale bar is 100 μm.

### IL-1β treatment induces MUC5B expression in control colonic organoids

As part of the inflammatory response to bacterial antigens, colonic epithelial cells secrete the pro-inflammatory cytokine IL-1β to recruit immune cells to the afflicted region of the gut (15). To determine whether IL-1β directly stimulates acidic mucin expression in the human colonic epithelium, differentiated control colonoids were treated with defined concentrations of recombinant IL-1β for 48 hrs. and sections were stained with Alcian blue. IL-1β treatment stimulated secretion of acidic mucins into the lumens of the colonic organoids (Fig. 4A). We next measured *MUC5B* transcripts and found that treatment with IL-1β (1 ng/ml) increased *MUC5B* expression (Fig. 4B). In addition, treatment of control organoids with IL-1β caused a dose-dependent increase in MUC5B proteins (Fig. 4C).

**Fig. 4.**
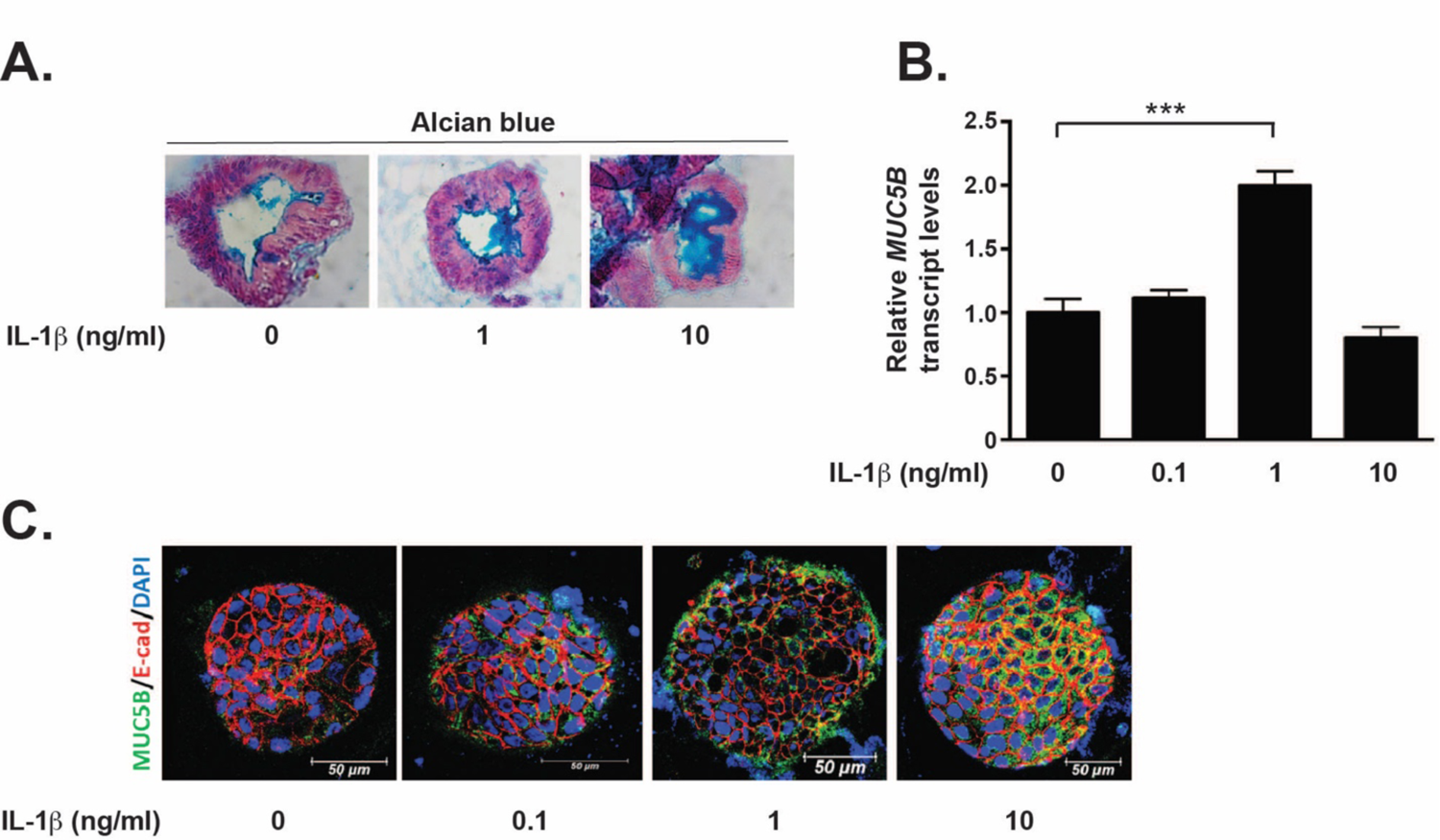
IL-1β treatment induces MUC5B expression in control colonic organoids. (A) Alcian blue staining of acidic mucins expressed in colonic organoids that were either untreated or treated with IL-1β (B) RT-qPCR analysis of *MUC5B* transcripts in untreated or IL-1β-treated colonic organoids (n=3). *MUC5B* expression levels are normalized to *GAPDH* and error bars are SEM, \*\*\**P* < 0.001. (C) Immunofluorescent analysis of control organoids that were untreated or treated with IL-1β and stained with DAPI or antibodies directed against MUC5B or E-cadherin.

### IL-1β treatment fails to induce MUC5B transcripts in UC organoids

We next determined whether the IL-1β-dependent increase of *MUC5B* transcripts in control organoids was also seen in UC organoids. We treated an independent set of control organoids (n=3) and UC organoids (n=4) with 1 ng/ml of IL-1β, and consistent with data presented in Fig. 4B, this caused an increase in *MUC5B* transcripts in control organoids (Fig. 5). However, treatment of UC organoids with this pro-inflammatory cytokine failed to induce *MUC5B* expression.

**Fig. 5.**
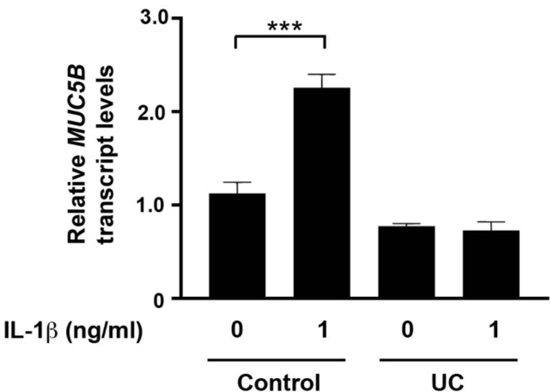
IL-1β treatment fails to induce *MUC5B* transcripts in UC organoids. RT-qPCR analysis of *MUC5B* transcripts in untreated or IL-1β-treated colonic organoids. Colonic organoids derived from healthy controls (n=3) and from UC tissues (n=4) were evaluated. *MUC5B* expression levels are normalized to *GAPDH* and error bars are SEM, \*\*\**P* < 0.001.

## DISCUSSION

Heavily-glycosylated mucin proteins are the primary constituents of a gelatinous mucus layer that coats intestinal epithelial cells and is in direct contact with the luminal environment (4–6). In the gut, MUC2 is the most abundant gel-forming mucin expressed and its protective function of the colonic epithelium is well-established (6–10). MUC5B is a second gel-forming mucin whose function has been primarily investigated in the lung (21–23). Although MUC5B is expressed in the colonic epithelium, its function in this tissue is largely unknown (11–14). In this study, we found that *MUC5B* transcripts are broadly expressed across various segments of the human colon. In addition, MUC5B protein expression was significantly reduced in inflamed UC colon tissues and in colonic organoids cultured from inflamed UC tissues. Moreover, we found that treating healthy colonic organoids with the pro-inflammatory cytokine, IL-1β, induced expression of *MUC5B* transcripts and proteins. IL-1β treatment failed to induce *MUC5B* transcripts in organoids derived from inflamed UC colonic segments. Together, these results demonstrate differential expression of MUC5B in healthy, non-IBD colonic versus UC colonic epithelia.

At the protein level, MUC5B is structurally most similar to MUC5AC (6, 7). Both proteins contain three von Willebrand D (VWD) domains at their amino-terminal ends, followed by a large central domain containing Pro-Thr-Ser (PTS) repeats and CysD domains, and a C-terminal domain with a fourth VWD, a von Willebrand C (VWC) domain, and a Cys-not (CK) domain. The major difference is the number of CysD domains in the central region with eleven present in MUC5AC and seven in MUC5B. Studies of pig trachea have found that the mucus in this tissue is arranged into bundles with MUC5B forming a core that is coated by MUC5AC (24). In this configuration, MUC5B forms strand-like structures with MUC5AC presenting a more diffuse staining pattern (24, 25). In addition, purified MUC5AC and MUC5B form unique macromolecular structures due to differences in multimeric organizations through their N-terminal regions (26). These findings demonstrate that MUC5B and MUC5AC contribute to the protective mucus layer in the lung through distinct mechanisms, but it remains to be determined whether these structural and functional arrangements are maintained in the gut.

*Muc5AC* deletion exacerbates disease phenotypes in the dextran-sodium sulfate (DSS) murine model of ulcerative colitis (27). Whereas *Muc5b*^-/-^ mice displayed chronic infection of bacteria in the lungs and subsequent inflammation that failed to fully resolve, whether these mice also displayed aberrant intestinal phenotypes was not reported (22). An in-depth analysis of intestinal architecture, the mucus bilayer, inflammatory cytokines and immune cell infiltrates is necessary to reveal a potential role for Muc5B in intestinal homeostasis. Moreover, although lung pathologies found in *Muc5B*^-/-^ mice could preclude assessment of Muc5b in the murine models of UC (23), testing whether *Muc5B*^+/-^ mice are more susceptible to colitis in such models is necessary to fully understand the role of Muc5B in the pathogenesis of UC.

IL-1β treatment has been shown to increase *MUC5B* transcripts in human airway epithelial cells by stimulating NF-kB binding to a defined NF-kB consensus sequence in the *MUC5B* proximal promoter (28). In addition, IL-1β has been shown to stimulate *MUC5B* transcripts and proteins in human bronchial epithelial cells by regulating activities of the SAM-pointed domain containing ETS transcription factor (SPEDF) and the endoplasmic reticulum stress sensor, ER to nucleus signaling 2 (ERN2) (29). While we have shown that IL-1β increases MUC5B expression in healthy human colonic organoids (Fig. 4B, C), whether these or other pathways are involved in this up-regulation remain to be determined. In addition, we found that although *MUC5B* transcripts were induced by a specific concentration of IL-1β, MUC5B protein levels demonstrated a dose-dependent increase in treated organoids (Fig. 4B, C). These results suggest that post-transcriptional and/or post-translational regulatory mechanisms are also involved in the IL-1β-dependent increase in MUC5B. That *MUC5B* transcripts are not induced by IL-1β in organoids derived from UC tissues (Fig. 5) indicates a defect in this regulatory response.

A prior study found that *MUC5B* transcripts were increased in UC patients versus controls (13), although a separate study indicated no difference in MUC5B protein levels in biopsies from control patients and those in various UC disease states (14). Our previous transcriptome analyses of surgically-resected colonic tissues of UC patients found that increased *MUC5B* transcripts correlated with a delayed time-to-surgery after UC diagnosis (12). While differences in study design and patient populations likely contribute to these disparate findings, our results in the current study indicates that IL-1β acutely stimulates *MUC5B* expression in healthy organoids, but not UC organoids. Based upon its expression in healthy colonic epithelium, it is therefore possible that the increased expression of MUC5B by this pro-inflammatory cytokine is part of a reparative program to heal the colonic epithelium. It follows that this response is lost in the colonic epithelium of UC patients which is subjected to chronic inflammation. However, we cannot at this time exclude the alternative that increased MUC5B in the mucinous layer is deleterious to the epithelium. More work is needed using mouse models of UC and genetically engineered human organoid models to discern between these two distinct possibilities.

In this present work, we clearly detect reduced MUC5B expression in inflamed UC tissues versus controls. Differences in patient populations, the difficulty in assessing cause versus affect in human studies and the lack of studies addressing intestinal *Muc5b* in murine models of colitis make it difficult to pinpoint the role of MUC5B in the pathogenesis of UC at this time.

However, our findings that IL-1β treatment induces MUC5B expression in healthy colonic organoids suggest that this contributes to restoration of the colonic epithelium in a pro-inflammatory environment. Clearly this model requires more experimental validation and a better understanding of how MUC5B affects the permeability of the colonic mucus bilayer, but if true, these results would suggest that strategies to restore MUC5B expression would be of therapeutic benefit to UC patients.

## CONCLUSION

Expression of the gel-forming mucin, MUC5B, is decreased in UC colonic epithelium. These results prompt the need for further investigation into the role of MUC5B in intestinal homeostasis and in the pathophysiology of UC.

## Acknowledgments

We would like to thank Sue Deiling and Leonard Harris for their efforts in supporting the Carlino Family IBCRD biobank and facilitating research of the banked human colonic tissues. We thank members of the Koltun and Yochum labs for helpful discussions throughout the course of this work. This project was funded, in part, under a grant with the Pennsylvania Department of Health using Tobacco CURE Funds. The Department specifically disclaims responsibility for any analyses, interpretations or conclusions. This research was also supported by the Peter & Marshia Carlino Fund for Inflammatory Bowel Disease Research.

## Conflicts of interest

No conflicts to declare.

## Authors’ contributions

W.D., G.S.Y.: study concept and design; W.D.: acquisition of data; W.D., G.S.Y.: analysis and interpretation of data; W.D., G.S.Y.: drafting of the manuscript; W.D., W.A.K., G.S.Y.: critical revision of the manuscript for important intellectual content; W.D.: statistical analysis; W.D., W.A.K., G.S.Y.: administrative, technical, or material support; G.S.Y.: study supervision. All authors approved the final version of the manuscript for publication

## Abbreviations

IBD: inflammatory bowel disease

IL-1β: interleukin 1 beta

MUC2: mucin 2

MUC5B: mucin 5B

RT-qPCR: reverse transcription quantitative PCR

UC: ulcerative colitis

---

**Supplemental Table 1:** Immunohistochemical analysis of MUC5B expression.

| Number | Sample ID | Score |
| --- | --- | --- |
| 1 | 19188 RCN | 0 |
| 2 | 18268 RCN | 2 |
| 3 | 17413 RCN | 3 |
| 4 | 18268 LCN | 2 |
| 5 | 17413 LCN | 3 |
| 6 | 19265 LCN | 3 |
| 7 | 20101 RCN | 3 |
| 8 | 19195 RCN | 2 |
| 9 | 20169 RCN | 3 |
| 10 | 20101 LCN | 3 |
| 11 | 16015 LCN | 3 |
| 12 | 20015 LCN | 3 |
| 13 | 19389 LCN | 2 |
|  | **MEAN** | **2.46** |
|  | **SD** | **0.88** |

| Number | Sample ID | Score |
| --- | --- | --- |
| 1 | 19378 RCM | 2 |
| 2 | 777315 RCM | 3 |
| 3 | 18250 RCS | 2 |
| 4 | 19378 LCS | 2 |
| 5 | 18465 LCS | 0 |
| 6 | 19131 RCM | 1 |
| 7 | 18315 RCM | 0 |
| 8 | 18226 RCM | 2 |
| 9 | 19297 RCS | 0 |
| 10 | 19021 RCS | 0 |
| 11 | 18295 RCS | 3 |
| 12 | 19006 LCM | 1 |
| 13 | 18201 LCM | 1 |
| 14 | 17261 LCM | 2 |
| 15 | 18020 LCM | 2 |
| 16 | 19297 LCS | 3 |
| 17 | 17156 LCS | 1 |
| 18 | 15238 LCS | 3 |
| 19 | 17245 LCS | 3 |
| 20 | 18310 LCS | 1 |
| 21 | 303011 LCS | 2 |
| 22 | 314011 LCS | 1 |
| 23 | 338011 LCM | 2 |
| 24 | 367011 LCS | 2 |
| 25 | 8882621 LCM | 3 |
| 26 | 8882811 LCM | 3 |
| 27 | 8882971 LCS | 2 |
| 28 | 8881711 LCM | 2 |
|  | **MEAN** | 1.750 |
|  | **SD** | 1.09 |

## Notes

### Competing Interest Statement

The authors have declared no competing interest.

